# Reward Network Activations of Win versus Loss in a Monetary Gambling Task

**DOI:** 10.1101/2025.03.20.644448

**Authors:** Chella Kamarajan, Babak A. Ardekani, Ashwini Pandey, Gayathri Pandey, Sivan Kinreich, Weipeng Kuang, Jacquelyn L. Meyers, Bernice Porjesz

## Abstract

Reward processing is a vital function for health and survival and is impaired in various psychiatric and neurological disorders. Using a monetary gambling task, the current study aims to elucidate neural substrates in the reward network underlying evaluation of win versus loss outcomes, and their association with behavioral characteristics, such as impulsivity and task performance, and neuropsychological functioning. Functional MRI was recorded in thirty healthy, male community volunteers (mean age = 27.4 years) while they performed a monetary gambling task in which they bet with either 10 or 50 tokens and received feedback of whether they won or lost the bet amount. Results showed that a set of key brain structures in the reward network, including putamen, caudate nucleus, superior and inferior parietal lobule, angular gyrus, and Rolandic operculum, had greater blood oxygenation level dependent (BOLD) signal during win relative to loss trials, and the BOLD signals in most of these regions were highly correlated with one another. Further, exploratory bivariate analyses between these reward related regions and behavioral and neuropsychological domains showed significant correlations with moderate effect sizes, including: (i) negative correlations between non-planning impulsivity and activations in putamen and caudate regions, (ii) positive correlations between risky bets and right putamen activation, (iii) negative correlations between safer bets and right putamen activation, (iv) a negative correlation between short-term memory capacity and right putamen activity, and (v) a negative correlation between poor planning skills and left inferior occipital cortex activation. These findings contribute to our understanding of the neural underpinnings of monetary reward processing and their relationships to aspects of behavior and cognitive function. Future studies may confirm these findings with larger samples of healthy controls and extend these findings by investigating various clinical groups with impaired reward processing.

## 1. Introduction

Reward processing is a key neurocognitive function essential for survival in most species. Understanding the mechanisms of reward processing is crucial as they are fundamental to human cognition and behavior, including motivation, learning, and decision-making [1]. Further, dysfunction in reward processing is linked to several psychiatric disorders such as addiction, depression, and obesity [2–4], making it critical to elucidate neural substrates and behavioral correlates of reward processing. While primary rewards (e.g., food, sex) and secondary rewards (e.g., money, tokens, and verbal reinforcements, such as appreciation) [5] are important, in humans the utmost importance is placed on monetary rewards, as they can buy most of the other rewards [6]. Most human studies investigating reward processing have often used monetary incentives as a proxy for primary rewards [7]. Further, neuroimaging studies have indicated that primary rewards may evoke similar neural responses in humans in response to more abstract, or secondary rewards such as monetary incentives [8,9]. Studies have also shown that gambling tasks are better suited to examine the neural substrates of monetary reward processing in humans than other paradigms [10].

Over the decades, studies on animals and humans have employed various neuroimaging methods to elucidate neural substrates of reward processing. In particular, many studies have used functional MRI (fMRI) to elucidate brain structures that are related to aspects of reward processing [11–17]. These studies have identified a set of subcortical (i.e., ventral tegmental area, nucleus accumbens, putamen, caudate, pallidum, amygdala, thalamus, and hippocampus) and cortical regions (i.e., insula, parahippocampal gyrus, cingulum, orbitofrontal cortex, angular gyrus, superior parietal lobule, inferior parietal lobule, and dorsolateral prefrontal cortex) that are activated during reward processing [18]. Studies have also found that individual variability in behavioral characteristics can affect reward processing and its neural substrates [6,10,19,20]. More specifically, psychological characteristics (e.g., impulsivity) [20,21], task performance (e.g., opting for risky vs. safer choices in tasks assessing reward processing) [20,22], and neuropsychological variables (e.g., executive functions, memory, etc.) [20,22] can modulate monetary reward processing. However, only a few studies have examined associations between psychological, behavioral and neuropsychological characteristics and neural substrates that underlie or modulate reward processing.

Impulsivity, a key factor known to modulate the reward processing mechanism [23,24], has three core aspects: 1) acting on the spur of the moment (motor impulsivity), 2) not focusing on the task at hand (attentional or cognitive impulsivity), and 3) lack of planning without adequate thinking (non-planning impulsivity) [25,26]. A previous fMRI study reported that individual differences in impulsivity accounted for variations in the reward-related blood oxygenation level dependent (BOLD) response in the ventral striatum and the orbitofrontal cortex [27]. Further, decreased activation of the ventral striatum and anterior cingulate during reward processing was shown to correlate with high impulsivity in those with alcohol use disorder [23]. These findings emphasize the need to confirm the association between impulsivity and BOLD activation of specific brain structures during reward processing, particularly while evaluating monetary outcomes.

fMRI studies have identified specific brain regions associated with task-related behaviors. For example, putamen activity was shown to be associated with the stimulus-action-reward association [28] and with reward sensitivity [29], while the midbrain activation was linked to efficiency in task performance [30]. However, the association between brain activation and participant’s performance style or strategies during monetary reward processing, such as choosing risky bets against safer options after previous losses, has not been adequately studied. Therefore, one of the aims of the current fMRI study is to examine this association during a monetary gambling task.

Lastly, studies have reported associations between cognitive abilities measured with neuropsychological tests and neural activation in specific brain structures during reward processing. For example, the basal ganglia structures, particularly the putamen, was found to modulate working memory in a delayed-response task of a reward paradigm [31], while putamen activity was associated with cognitive functions in general [32] and learning and memory in particular [33,34]. Further, executive functions such as planning and problem solving may also be inherently associated with reward processing and related neural activation [35]. Although brain-behavior associations in the context of reward processing have been examined by some studies in an isolated fashion, no study has investigated multiple behavioral and neuropsychological domains on the same participants.

Therefore, the current study aims to understand associations among brain regions involved in reward processing networks and elucidate neural substrates underlying evaluation of win versus loss outcomes using a monetary gambling task in a group of healthy participants. Furthermore, the current study will investigate possible associations of activations of brain reward structures with key aspects of behavior and cognitive function (impulsivity, task performance, and neuropsychological measures).

## 2. Methods

### 2.1. Sample

The sample consisted of 30 healthy, young male participants (ages 19-38, mean=27.4 years) who were recruited from the community. The exclusion criteria were (i) diagnosis of major psychiatric disorder or substance use disorders, (ii) hearing/visual impairment, (iii) history of head injury, and (iv) cognitive deficits or a score of <24 on the mini-mental state examination (MMSE). Most of the participants (28 out of 30) in the study were right-handed, and their education ranged from 12-20 years with a mean of 15.8 years. Behavioral and neuropsychological data were collected at the SUNY Downstate Health Sciences University. The structural and functional MRI data were acquired at the Nathan Kline Institute (NKI) for Psychiatric Research. Informed consent was obtained from the participants, and the research protocol was approved by the Institutional Review Boards of both centers (IRB approval ID: SUNY–266893; NKI–212263).

### 2.2. The Monetary Gambling Task (MGT)

The Monetary Gambling Task (MGT), as illustrated in **Figure 1**, consisted of 240 trials. The duration of each trial was 2.5 seconds, and it took about 10 minutes to complete the task. Each trial consisted of two stimulus presentations: (a) a pie stimulus presenting the chance of winning (75%, 50%, or 25%) for the duration of 1.5 s, during which the participant was instructed to select a bet amount of either 10 tokens, by pressing button 1, or 50 tokens, by pressing button 2, on a button response unit with their right hand; and (b) a feedback stimulus for the duration of 1.0 s with a text indicating whether the participant won or lost the bet amount. Thus, six types of trials were presented randomly, irrespective of the bet amount (see **Table 1**): (1) chance of winning = 50%; outcome = win; number of trials = 40; (2) chance of winning = 50%; outcome = loss; number of trials = 40; (3) chance of winning = 75%; outcome = win; number of trials = 60; (4) chance of winning = 75%; outcome = loss; number of trials = 20; (5) chance of winning = 25%; outcome = win; number of trials = 20; and (6) chance of winning = 25%; outcome = loss; number of trials = 60. The participants were not made aware of the trial types and sequence. If the participant failed to make a bet within 1.5 s after the presentation of the pie stimulus, a feedback stimulus “No bet made!” would appear on the screen. The total amount won or lost by the participant was displayed at the end of the task.

**Figure 1.**
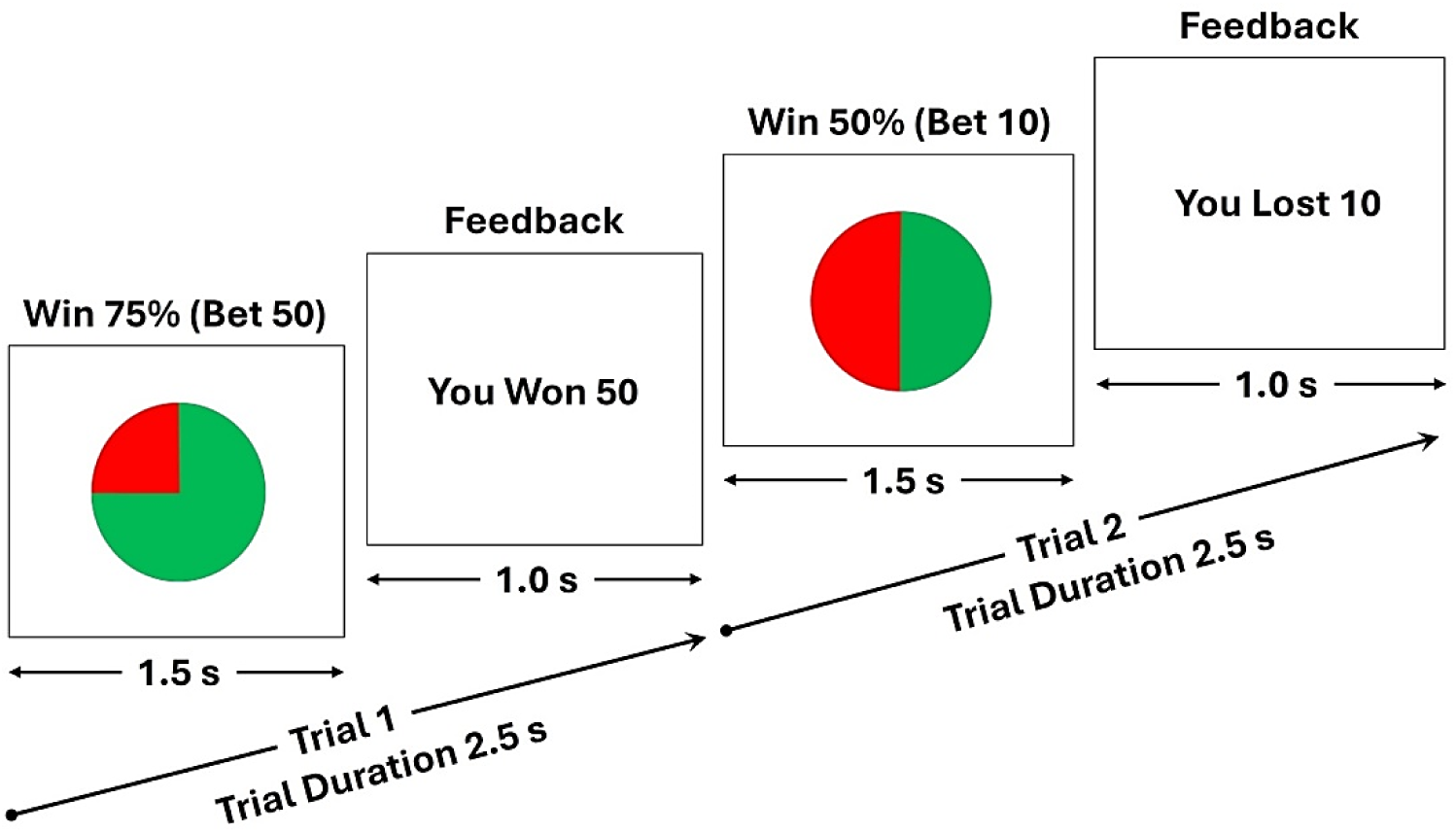
The schematic diagram of the monetary gambling task, showing two random trials: (i) a trial showing a pie stimulus (duration = 1.5 s) with the winning chance of 75% for which a participant bet with 50 tokens and won the bet amount as displayed in the feedback stimulus (duration = 1.0 s) (Trial 1); and (ii) another trial showing the pie stimulus with the winning chance of 50% for which the participant bet with 10 tokens and lost the bet amount as displayed in the feedback stimulus (Trial 2). The task consisted of 240 trials and the length of each trial was 2.5 seconds.

**Table 1.** Various trial types used in the monetary gambling task.

| **Trial Type** | **Chance of winning** | **Outcome** | **Number of trials** |
| --- | --- | --- | --- |
| 1 | 50% | Win | 40 |
| 2 | 50% | Loss | 40 |
| 3 | 75% | Win | 60 |
| 4 | 75% | Loss | 20 |
| 5 | 25% | Win | 20 |
| 6 | 25% | Loss | 60 |

### 2.3. Behavioral Scores Extracted from Task Performance

The list of behavioral scores that were computed for the participants based on their performance of the monetary gambling task is shown in **Table 2**. These scores included: (i) total number of tokens won or lost at the end of the task; (ii) number of trials with bet amount 50 when the outcome/feedback was “Loss” for the previous one, two, and three trials, respectively, suggesting potential risky behavior; (iii) number of trials with bet amount 10 when the outcome/feedback was “Loss” for the previous one, two, and three trials, respectively, suggesting potential safe behavior; (iv) number of trials with bet amount 50 when the net outcome was “Loss” for the previous two and three trials, respectively, indicating risky behavior; and (v) number of trials with bet amount 10 when the net outcome was “Loss” for the previous two and three trials, respectively, indicating safe behavior. The term “net outcome” for two or more consecutive trials refers to the resulting number of tokens lost or won during those trials. For instance, if the outcomes during the previous two trials were a win of 10 and a loss of 50, the net outcome for those two trials would be a loss of 40 tokens.

**Table 2.** Behavioral scores extracted from the performance data of the monetary gambling task.

| **Variable Name** | **Variable Description** |
| --- | --- |
| Net_Outcome | Total tokens won or lost at the end of the task |
| Bet50_Priv1Loss | Number of bets with 50 tokens after a loss during the previous trial |
| Bet10_Priv1Loss | Number of bets with 10 tokens after a loss during the previous trial |
| Bet50_Priv2Loss | Number of bets with 50 tokens after two consecutive losses during the previous trials |
| Bet10_Priv2Loss | Number of bets with 10 tokens after two consecutive losses during the previous trials |
| Bet50_Priv3Loss | Number of bets with 50 tokens after three consecutive losses during the previous trials |
| Bet10_Priv3Loss | Number of bets with 10 tokens after three consecutive losses during the previous trials |
| Bet50_Priv2NetLoss | Number of bets with 50 tokens after the net outcome of loss during the previous two trials |
| Bet10_Priv2NetLoss | Number of bets with 10 tokens after the net outcome of loss during the previous two trials |
| Bet50_Priv3NetLoss | Number of bets with 50 tokens after the net outcome of loss during the previous three trials |
| Bet10_Priv3NetLoss | Number of bets with 10 tokens after the net outcome of loss during the previous three trials |

### 2.4. Neuroimaging Protocol

#### 2.4.1. Structural and Functional MRI Data Collection

Both structural and functional MRI data were collected using a 3.0 Tesla Siemens Trio scanner (Erlangen, Germany). BOLD fMRI scans were acquired using a T2*-weighted gradient echo single-shot echo-planar imaging (EPI) sequence with these acquisition parameters: number of slices=36; voxel size=(2.5×2.5×3.5) mm^3^; FOV=240 mm; TR=2500 ms; TE=30 ms; flip angle=80°; parallelization factor=2; acquisition time=2.5 s per volume; and number of volumes=240. The sequence was carefully optimized to minimize the effects of magnetic susceptibility inhomogeneities (such as distortions and signal dropouts), as well as the effects of mechanical vibrations, which elevate Nyquist ghosting levels. In addition, a magnetization-prepared rapid gradient-echo (MPRAGE) high-resolution three-dimensional T1-weighted structural sequence was collected to be used as anatomical reference for the fMRI data and for the non-linear registration of imaging data between subjects. The sequence parameters for the MPRAGE were: TR = 2500 ms; TE = 3.5 ms; TI = 1200 ms; flip angle = 8°; voxel size = 1 × 1 × 1 mm^3^; matrix size = 256 × 256 × 192; FOV = 256 mm; and number of averages = 1.

#### 2.4.2. Image Processing of the fMRI data

Processing of the imaging data included the following stages. Within each subject, the MPRAGE and fMRI volumes were registered using the intra-subject inter-modality linear registration module [36] of the Automatic Registration Toolbox (ART; www.nitrc.org/projects/art). The *brainwash* program within the ART toolbox was used for skull-stripping the MPRAGE volumes. To correct for small subject motion during fMRI acquisitions, motion detection and correction was performed using the *3dvolreg* module of the AFNI software package [37]. To correct for the geometric distortions of the fMRI images due to magnetic susceptibility differences in the head, particularly at brain/air interfaces, we used the non-linear registration module of the ART [38]. The skull-stripped MPRAGE images from all subjects were non-linearly registered to a study-specific population template using ART’s non-linear registration algorithm *3dwarper*, which is one of the most accurate inter-subject registration methods available [39]. The population template was formed using an iterative method [40]. The motion-corrected fMRI time series were detrended using PCA [41]. Finally, fMRI data from all subjects were normalized to a standard space using the image registration steps outlined above, which were mathematically combined into a single transformation and used in re-sampling the fMRI data.

#### 2.4.3. Extraction of BOLD Activation Clusters for the Win-Loss Contrast

We used a Sparse Principal Component Analysis (sPCA) method [42] to identify activation clusters from the data matrix containing 7050 voxels that were detected to be activated during the task conditions. The sPCA is a novel and efficient method to identify activation clusters. While the components from the sPCA have a natural ordering according to their variance similar to the regular PCA, the sPCA performed better in separating the noise from the signal and was found to be more flexible, less committed, and easier to interpret [42,43], The sPCA extracts relatively a minimal number of non-zero components by using the LASSO regression technique, which by driving some loadings to exactly zero and adjusting the other components to approximate the properties of the PCA [42]. We used the top clusters with a size of 100 voxels or more for statistical analyses (see **Table 3**). Activation value for each cluster was determined by taking the mean from all of its voxels. The anatomical label for the clusters, based on the MNI coordinates of their centroids, were extracted using the automated anatomical labeling (AAL) method [44], as provided by the R-package *label4MRI* (https://github.com/yunshiuan/label4MRI) (see **Table 3** and **Figure 2**).

**Figure 2.**
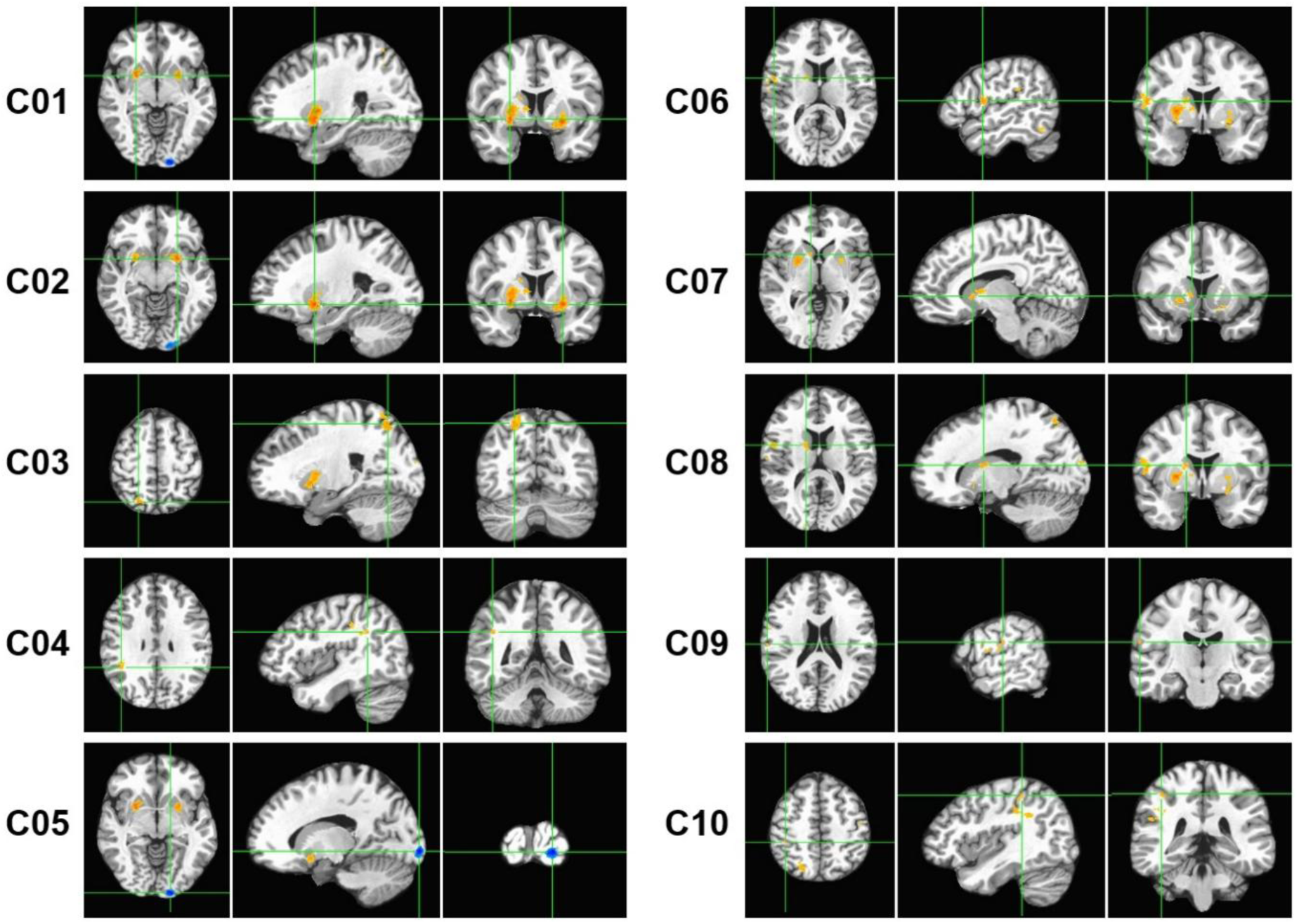
The 10 fMRI activation clusters (C01–C10) with 100 or more voxels that were extracted from the Win-Loss contrast of the monetary gambling task. The centroid of each cluster is shown with green crosshair lines for the axial (left panels), sagittal (middle panels), and coronal (right panels) views. The activated voxels are highlighted in orange/red (Win > Loss) or cyan/blue (Loss > Win).

**Table 3:** The fMRI activation clusters for the Win-Loss contrast, which had 100 or more voxels. The number of voxels, anatomical region, Brodmann area, MNI, and voxel coordinates are shown for each cluster.

| Cluster Number | Size (voxels) | Anatomical Region | Abbreviation | Direction of Activation | Brodmann Area | MNI Coordinates |  |  |
| --- | --- | --- | --- | --- | --- | --- | --- | --- |
|  |  |  |  |  |  | x | y | z |
| 1 | 1781 | R. Putamen | R. Ptm | Win > Loss | 49 | 27 | 5 | -6 |
| 2 | 1426 | L. Putamen | L. Ptm | Win > Loss | 49 | -24 | 5 | -9 |
| 3 | 878 | R. Superior Parietal Lobule | R. SPL | Win > Loss | 7 | 23 | -68 | 56 |
| 4 | 663 | R. Angular Gyrus | R. AnGy | Win > Loss | 39 | 44 | -47 | 30 |
| 5 | 640 | L. Inferior Occipital Cortex | L. IOC | Loss > Win | 18 | -15 | -100 | -6 |
| 6 | 444 | R. Rolandic Operculum | R. RoOp | Win > Loss | 6 | 56 | 2 | 12 |
| 7 | 333 | R. Caudate (anterior-inferior) | R. Cdt (A-I) | Win > Loss | 48 | 11 | 12 | 0 |
| 8 | 239 | R. Caudate (posterior-superior) | R. Cdt (P-S) | Win > Loss | 48 | 17 | 1 | 14 |
| 9 | 100 | R. Supramarginal Gyrus | R. SMG | Win > Loss | 40 | 63 | -18 | 20 |
| 10 | 100 | R. Inferior Parietal Lobule | R. IPL | Win > Loss | 40 | 42 | -37 | 51 |

L – Left hemisphere; R – Right hemisphere

### 2.5. Assessment of Impulsivity

The participants were administered the Barratt Impulsiveness Scale—Version 11 (BIS-11) [26], a 30-item self-administered tool that assessed motor impulsivity (BIS_MI), non-planning (BIS_NP), attentional impulsivity (BIS_AI), and total score (BIS_Tot). BIS-11 has been widely used to assess impulsivity and its biological, psychological, and behavioral correlates [45]. Attentional impulsivity refers to an inability to focus attention or concentrate on the job at hand, motor impulsivity is the tendency to act on the spur of the moment without thinking, while non-planning impulsivity is conceptualized as a lack of forethought or planning for executing a task [46].

### 2.6. Neuropsychological Assessment

Computerized adaptations of the Tower of London Test (TOL) [47] and the Visual Span Test (VST) [48,49] were administered using the Colorado assessment tests for cognitive and neuropsychological assessment [50], as described previously [51]. Details of these tests are summarized below.

#### 2.6.1. Tower of London Test (TOL)

The planning and problem-solving ability of the executive functions were assessed using the TOL in which participants solved a set of puzzles with graded difficulty levels by arranging the color beads one at a time from a starting position to a desired goal position in as few moves as possible. The test consisted of 3 puzzle types with 3, 4, and 5 colored beads placed on the same number of pegs, with 7 problems/trials per type and a total of 21 trials. Five performance measures from the summation of all puzzle types were used in the analysis: (i) excess moves (additional moves beyond the minimum moves required to solve the puzzle); (ii) average pickup time (initial thinking/planning time spent until picking up the first bead to solve the puzzle); (iii) average total time (total thinking/planning time to solve the problem in each puzzle type); (iv) total trial time (total performance/execution time spent on all trials within each puzzle type); and (v) average trial time (mean performance/execution time across trials per puzzle type).

#### 2.6.2. Visual Span Test (VST)

The VST was used to assess visuospatial memory span from the forward condition and working memory from the backward condition. In this test, 8 randomly arranged squares were displayed on the screen, and 2–8 squares flashed in a predetermined sequence depending on the span level being assessed. Each span level was administered twice, with a total of 14 trials in each condition. During the forward condition, subjects were required to repeat the sequence in the same order via mouse clicks on the squares. In the backward condition, subjects were required to repeat the sequence in reverse order (starting from the last square). Four performance measures were collected during forward and backward conditions (with a total of 8 scores): (i) total correct scores (total number of correctly performed trials); (ii) span (maximum sequence length achieved); (iii) total average time (sum of the mean time taken across all trials performed); and (iv) total correct average time (sum of the mean time taken across all trials correctly performed).

### 2.7. Statistical Analyses

All statistical analyses were performed using the R packages [52]. Pearson bivariate correlations were performed to identify significant relationships across different fMRI activations clusters. We also performed an explorative correlational analysis to test the associations between the fMRI activation clusters and the behavioral/cognitive variables. In order to avoid the Type 2 error [53,54] due to the small sample size, multiple testing corrections were not performed for the explorative correlational analysis, and the strength of association was determined based on the magnitude of correlation coefficient as a metric of effect size [55].

## 3. Results

### 3.1. The fMRI Activation Clusters for the Win-Loss Contrast

As shown in **Table 3** and **Figure 2**, the BOLD activation difference between the gambling outcomes (Win-Loss) elicited 10 regions (i.e., clusters) with 100 or more voxels showing activations: (i) Right Putamen, (ii) Left Putamen, (iii) Right Superior Parietal Lobule, (iv) Right Angular Gyrus, (v) Left Inferior Occipital Cortex, (vi) Right Rolandic Operculum, (vii) Right Caudate (anterior-inferior), (viii) Right Caudate (posterior-superior), (ix) Right Supramarginal Gyrus, and (x) Right Inferior Parietal Lobule. Eight of the ten clusters were on the right side, with only two clusters on the left, namely, left putamen and left inferior occipital cortex. Nine of the ten clusters showed higher activation during win relative to loss (orange/red blobs in **Figure 2**), with only the left inferior occipital cortex (cluster # 5) showing higher activation during loss (cyan/blue blobs in **Figure 2**). Overall, eight clusters represented the right hemisphere, and two clusters were of the left hemisphere. Putamen showed bilateral activations represented by clusters 1 & 2. Right Caudate was represented by two separate clusters in different locations (i.e., cluster 7 was at the anterior-inferior location and cluster 8 was at the posterior-superior location. Interestingly, three of the cortical clusters (i.e., clusters 4, 9, and 10) represented inferior parietal lobule, and two of them (clusters 9 & 10) represented the same Brodmann area (BA 40) although cluster 10 (R. IPL) was more medial, posterior, and superior to the anatomical location of cluster 9 (R. SMG).

Correlations among the fMRI activation clusters are shown in **Figure 3**. All clusters except cluster 5 (L. IOC) showed significant positive correlations with other clusters, even after correcting for multiple testing. Bonferroni-adjusted significant correlations were between (i) cluster 1 (R. Ptm) and cluster 2 (L. Ptm); (ii) cluster 1 (R. Ptm) and cluster 7 (R. Cdt A-I); (iii) cluster 1 (R. Ptm) and cluster 8 (R. Cdt P-S); (iv) cluster 2 (L. Ptm) and cluster 3 (R. SPL); (v) cluster 2 (L. Ptm) and cluster 4 (R. AnGy); (vi) cluster 3 (R. SPL) and cluster 4 (R. AnGy); (vii) cluster 4 (R. AnGy) and cluster 10 (R. IPL); (viii) cluster 6 (R. RoOp) and cluster 9 (R. SMG); and (ix) cluster 7 (R. Cdt A-I) and cluster 8 (R. Cdt P-S). Overall, clusters 1, 2, and 4 had three significant correlations each, followed by clusters 3, 7, and 8 had two correlations each, and clusters 6 and 9 had a single correlation with each other.

**Figure 3.**
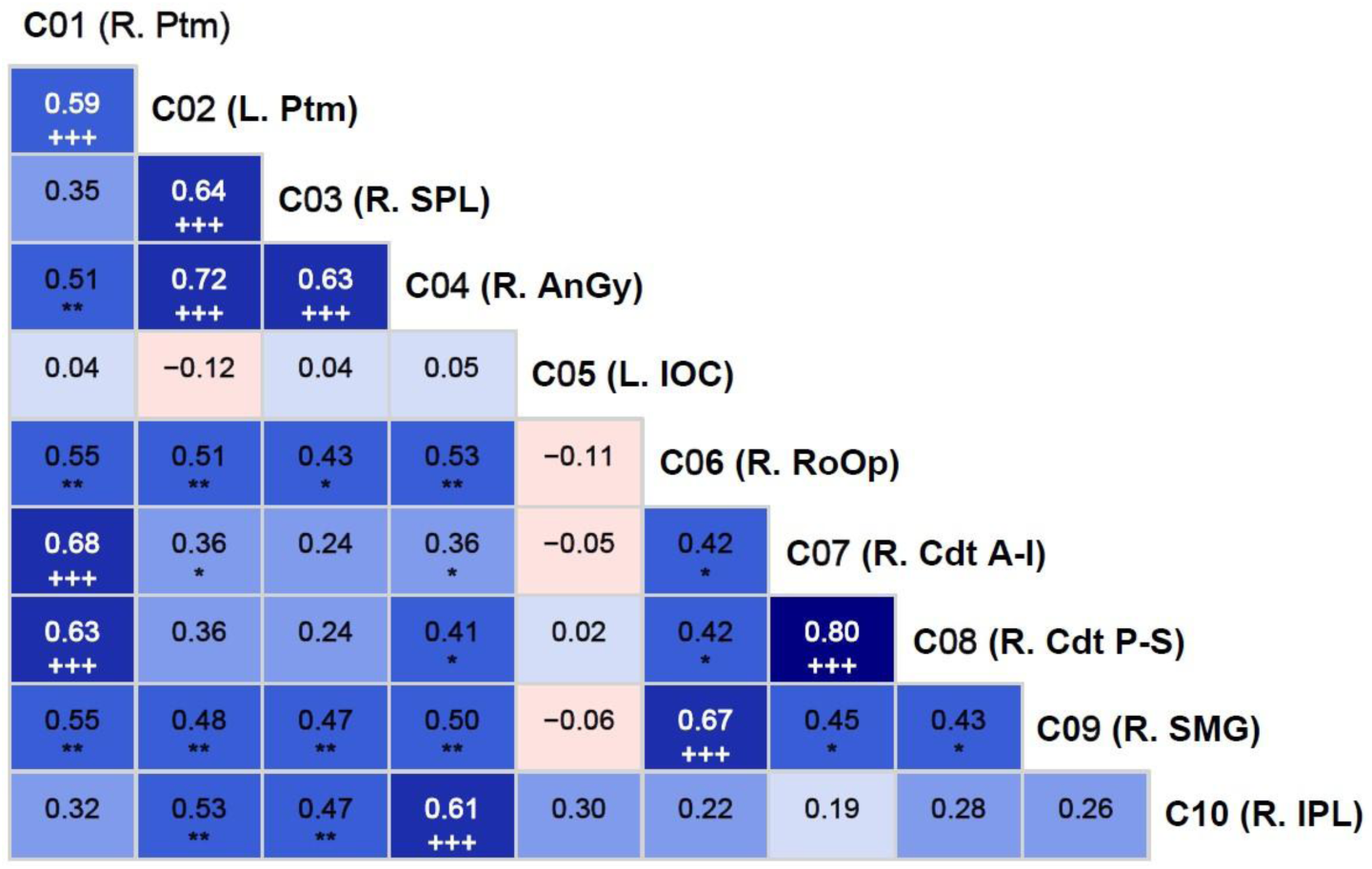
Pearson bivariate correlations among the fMRI activation clusters (C01-C10). The correlation coefficient (number) and the level of significance (asterisks and plus signs) are provided within each cell. The asterisks (*p<0.05; **p<0.01; ***p<0.001) represent significance before the Bonferroni corrections, while the plus sign (+++) indicates those that survived Bonferroni correction. The cyan/blue shades represent positive correlations, and the pink shades indicate negative correlations.

### 3.2. Correlations Between the fMRI Activation Clusters and Other Variables

Pearson bivariate correlations between the fMRI clusters (C01-C10) and other variable sets (demographic variables, impulsivity scores, task performance measures, and neuropsychological performance) are shown in **Table 4**. While some correlations in each variable set (except in demographic set) were significant and with moderate-high effect sizes ranging from 0.3617 to 0.5603 [55], none of these correlations survived multiple testing corrections due to small sample size. When sample size is small, as in our case, it may be better to rely on the effect size of the correlations rather than on multiple testing corrections on the significant correlations [53,54]. Therefore, we have opted to identify significant correlations based on their effect sizes in order to avoid Type 2 error (see *Section 2.7*). These significant correlations within each variable set are listed below:

**Table 4.** Pearson bivariate correlations between the fMRI clusters (C01-C10) and other variable sets, such as demographic variables, impulsivity scores, task performance, and neuropsychological scores [see the *Methods* section for the details of these variables].

| Variable Set | Variable | C01 | C02 | C03 | C04 | C05 | C06 | C07 | C08 | C09 | C10 |
| --- | --- | --- | --- | --- | --- | --- | --- | --- | --- | --- | --- |
| Demographic variables | Age | -0.1686 | -0.0885 | -0.3225 | -0.0864 | 0.0366 | -0.2015 | 0.0519 | 0.1424 | 0.0452 | 0.1024 |
|  | Education | -0.1406 | -0.1360 | -0.1668 | -0.1591 | 0.2654 | -0.1568 | -0.1810 | 0.0538 | -0.1594 | 0.1576 |
| Impulsivity scores | BIS_NP | **-0.3844\*** | **-0.4057\*** | -0.3534 | -0.3291 | -0.2510 | -0.2459 | **-0.4072** | **-0.5603\*\*** | -0.2798 | -0.3155 |
|  | BIS_MI | -0.0127 | -0.0581 | **-0.3885\*** | -0.0754 | -0.2758 | -0.2030 | -0.1888 | -0.0924 | -0.0088 | 0.1499 |
|  | BIS_AI | -0.0844 | -0.3143 | -0.1279 | -0.0107 | -0.0940 | -0.1215 | -0.1463 | -0.3095 | -0.1738 | -0.1755 |
|  | BIS_Tot | -0.2422 | -0.3562 | **-0.3851\*** | -0.2091 | -0.2746 | -0.2543 | -0.3446 | **-0.4504\*** | -0.2178 | -0.1770 |
| Task performance | Net_Outcome | -0.1250 | -0.2576 | 0.1526 | -0.2325 | 0.1594 | -0.3039 | 0.0517 | -0.0319 | -0.2912 | 0.0202 |
|  | Bet50_Priv1Loss | **0.3700\*** | 0.0915 | 0.1586 | 0.1642 | 0.1348 | **0.3617\*** | 0.1382 | 0.0521 | 0.0873 | 0.0799 |
|  | Bet10_Priv1Loss | -0.2937 | -0.1051 | -0.0691 | -0.0909 | -0.2319 | -0.1873 | -0.1078 | 0.0133 | -0.1279 | -0.0318 |
|  | Bet50_Priv2Loss | **0.3754\*** | 0.1812 | 0.2366 | 0.2337 | -0.0326 | **0.3896\*** | 0.1347 | 0.0513 | 0.1883 | 0.1301 |
|  | Bet10_Priv2Loss | **-0.3903\*** | -0.1510 | -0.0632 | -0.0695 | -0.1868 | -0.2537 | -0.1929 | -0.0736 | -0.2222 | -0.0041 |
|  | Bet50_Priv3Loss | 0.2540 | 0.1339 | 0.2415 | 0.2929 | -0.0371 | 0.3139 | 0.1563 | 0.1143 | 0.2803 | 0.1552 |
|  | Bet10_Priv3Loss | **-0.3943\*** | -0.0221 | 0.0013 | -0.0020 | -0.1044 | -0.2551 | -0.2749 | -0.1478 | -0.3456 | 0.1476 |
|  | Bet50_Priv2NetLoss | 0.2952 | 0.0196 | -0.0084 | 0.0484 | 0.0049 | 0.3052 | 0.0501 | -0.0094 | 0.1951 | 0.0100 |
|  | Bet10_Priv2NetLoss | -0.2676 | -0.0495 | -0.0523 | -0.0499 | -0.2094 | -0.1299 | -0.0098 | 0.1107 | -0.0841 | 0.0173 |
|  | Bet50_Priv3NetLoss | 0.3246 | 0.0931 | 0.0230 | 0.1136 | 0.0553 | 0.3604 | 0.0022 | -0.0122 | 0.1900 | 0.0138 |
|  | Bet10_Priv3NetLoss | -0.1418 | 0.0355 | 0.0016 | 0.0317 | -0.2043 | -0.0929 | 0.0215 | 0.1228 | -0.0544 | 0.0315 |
| Neuropsychological scores | TOL_ExcMovMade_All | -0.0468 | 0.0523 | -0.0938 | -0.1615 | **-0.3927\*** | 0.0679 | -0.0179 | -0.1394 | -0.0035 | -0.1466 |
|  | TOL_AvgPicTime_All | 0.0411 | 0.1848 | 0.2992 | 0.1470 | 0.2767 | -0.1186 | 0.1002 | 0.1636 | 0.1843 | 0.0613 |
|  | TOL_AvgTotTime_All | 0.0533 | 0.1626 | 0.1840 | 0.0473 | 0.2300 | -0.2029 | 0.0916 | 0.1049 | 0.0622 | -0.0133 |
|  | TOL_TotTrlTime_All | 0.0514 | 0.1852 | 0.1691 | 0.0168 | 0.1580 | -0.1979 | 0.0893 | 0.0756 | 0.0297 | -0.0447 |
|  | TOL_AvgTrlTime_All | 0.0514 | 0.1852 | 0.1691 | 0.0168 | 0.1580 | -0.1979 | 0.0894 | 0.0756 | 0.0297 | -0.0447 |
|  | VST_TotCor_Fw | **-0.3981\*** | -0.2994 | -0.0212 | -0.1768 | 0.1453 | -0.1415 | -0.1193 | -0.0483 | 0.0324 | -0.0508 |
|  | VST_Span_Fw | -0.3102 | -0.2193 | 0.0114 | 0.0489 | 0.0826 | 0.0098 | -0.0375 | 0.0441 | 0.1885 | 0.0089 |
|  | VST_TotAvgTime_Fw | -0.1477 | -0.0876 | 0.0234 | 0.0592 | 0.1884 | -0.0288 | 0.1645 | 0.1863 | 0.2613 | -0.0152 |
|  | VST_TotCorAvgTime_Fw | -0.1073 | 0.0129 | 0.0125 | 0.0592 | 0.1609 | -0.0083 | 0.1710 | 0.1496 | 0.2279 | 0.0012 |
|  | VST_TotCor_Bw | 0.0308 | 0.0717 | 0.2810 | 0.0232 | -0.0563 | 0.0489 | 0.2737 | 0.2256 | 0.2482 | -0.0321 |
|  | VST_Span_Bw | 0.0492 | -0.0505 | 0.1533 | -0.1092 | -0.0339 | -0.0772 | 0.2028 | 0.1715 | 0.1973 | -0.0333 |
|  | VST_TotAvgTime_Bw | 0.1166 | 0.0429 | 0.1718 | -0.0298 | -0.2722 | 0.0345 | 0.2830 | 0.2251 | 0.2971 | -0.1202 |
|  | VST_TotCorAvgTime_Bw | 0.1548 | 0.0176 | 0.2422 | -0.0960 | -0.1317 | -0.0426 | 0.2714 | 0.1209 | 0.3142 | -0.0829 |

\* p<0.05; \*\*p<0.01

#### Impulsivity

i. Negative correlation of BIS non-planning with fMRI activation cluster 1 (R. Ptm; r=-0.3844, p<0.05), cluster 2 (L. Ptm; r=-0.4057, p<0.05), cluster 7 (R. Cdt A-I; r=-0.4073, p<0.05), and cluster 8 (R. Cdt P-S; r=-0.5603, p<0.01); and
ii. Negative correlation of BIS motor impulsivity with cluster 3 (R. SPL; r=-0.3885, p<0.05);
iii. Negative correlation of BIS total impulsivity with cluster 3 (R. SPL; r=-0.3851, p<0.05) and cluster 8 (R. Cdt P-S; r=-0.4504).

#### Task Performance

i. Positive correlations between the number of bets with 50 after a loss during the previous trial with the fMRI activation cluster 1 (R. Ptm; r=0.3700, p<0.05) and cluster 6 (R. RoOp; r=0.3617, p<0.05);
ii. Positive correlations between the number of bets with 50 after two consecutive losses during the previous trials with the fMRI activation cluster 1 (R. Ptm; r=0.3754, p<0.05) and cluster 6 (R. RoOp; r=0.3896, p<0.05); and
iii. Negative correlations of the fMRI activation cluster 1 (R. Ptm) with the number of bets with 10 tokens after consecutively losing during the previous two trials (r=-0.3903, p<0.05) as well as with the number of bets with 10 tokens after consecutively losing during the previous three trials (r=-0.3943, p<0.05).

## 4. Discussion

The current study aimed to elucidate neural substrates of monetary reward outcomes and their association with behavioral and cognitive features. Ten BOLD activation clusters were identified for the win-loss contrast (see **Table 3** and **Figure 2**), which included bilateral putamen, right caudate nucleus, right superior and inferior parietal lobule, right angular gyrus, and right rolandic operculum. These anatomical regions showed greater activation during the win condition relative to the loss condition. It was found that all clusters except cluster 5 (left inferior occipital cortex) showed significant positive correlations with other clusters, suggesting that these brain regions were activated together during reward processing. Further, exploratory bivariate correlations with moderate effect sizes suggested possible associations between these reward regions and some behavioral and cognitive characteristics, including: (i) negative correlations between non-planning impulsivity and activations in putamen and caudate regions, (ii) positive correlations between risky bets and right putamen activation, (iii) negative correlations between safer bets and right putamen activation, (iv) a negative correlation between short-term memory capacity and right putamen activity, and (v) a negative correlation between poor planning skills and left inferior occipital cortex activation.

### 4.1. Neural Substrates of the Win-Loss Contrast

#### 4.1.1. The regions activated during reward processing

The current study identified ten BOLD activation clusters for the win-loss contrast (see **Table 3** and **Figure 2**), which include (i) Right Putamen, (ii) Left Putamen, (iii) Right Superior Parietal Lobule, Right Angular Gyrus, (v) Left Inferior Occipital Cortex, (vi) Right Rolandic Operculum, (vii) Right Caudate (anterior-inferior), (viii) Right Caudate (posterior-superior), (ix) Right Supramarginal Gyrus, and (x) Right Inferior Parietal Lobule. It is worth noting that most of these regions were part of the reward network as reported by the previous studies [13,14]. Our study has identified brain structures of the dorsal striatum, such as putamen (clusters 1 and 2) and caudate nucleus (clusters 7 and 8), which are the core structures of the reward network [13]. According to Arsalidou et al. [14], a common reward processing circuit is composed of basal ganglia nuclei such as the caudate, putamen, and globus pallidus, and these nuclei represent a basic subcortical structure that subserve reward processes irrespective of the reward outcome type or contextual factors associated with the rewards. Anatomically, both Putamen and caudate project to the globus pallidus, which in turn has projections with the thalamus (Ikemoto et al. 2015). Broadly, the mesocorticolimbic reward system includes dopaminergic projections from the ventral tegmental area to both nucleus accumbens and dorsal striatum (i.e., caudate and putamen) as well as orbital frontal cortex (OFC), medial prefrontal cortex (mPFC), and amygdala [56]. Although our study did not implicate the ventral striatum (i.e., nucleus accumbens), putamen and caudate structures are more involved in the monetary reward processing than other basal ganglia structures [14].

In terms of laterality, eight of the 10 clusters represented the right hemisphere, except the two left-hemisphere clusters. A meta-analysis of fMRI studies on reward processing indicated that monetary rewards activated all the basal ganglia nuclei bilaterally with the exception of the lateral globus pallidus [14]. In our study, while the putamen was involved bilaterally, only the right caudate was implicated for the win-loss contrast, along with other right hemisphere regions, such as the superior parietal lobule, angular gyrus, supramarginal gyrus, and rolandic operculum. The superior parietal lobule is involved in visual attention, spatial perception, visuomotor functions, spatial reasoning, and visual working memory [57], which are important elements during the evaluation of rewards and risk while performing the visual monetary gambling task as used in our study. On the other hand, regions of the inferior parietal lobule (cluster 10), such as angular gyrus (cluster 4) and supramarginal gyrus (cluster 9), are known to be involved in language ability, future planning, problem-solving, calculations, and other complex mental operations [58], some of which are essential while processing monetary reward stimuli. Besides, other neuroimaging studies have implicated the inferior parietal lobule to evaluate the possible motor significance of sensory stimuli [59] and perceptually based decisions and prospective action judgment [60], the functions that are also essential during the performance of gambling task. The right rolandic operculum (cluster 6) was found to be associated with affective evaluation and depression [61], possibly with loss outcomes during the performance of a gambling task. Lastly, the left inferior occipital cortex (cluster 5; Brodmann area 18) showed higher activation during loss compared to win outcomes. This secondary visual association cortex is known to be involved in visual processing of color, motion, and depth perception [62], functions that are practically imperative to perform the gambling task that contains the processing of visual stimuli presented on a computer screen. Overall, most of the brain regions activated for the win-loss contrast in our study are key regions of the reward circuitry and are consistent with the previous findings on monetary reward processing.

#### 4.1.2. Correlations Across the fMRI Activation Clusters

One of the sub-aims of the study is to determine if the reward-related activation clusters are correlated or connected with one another. As shown in **Figure 3**, all clusters except cluster 5 (L. IOC) showed significant positive correlations with one or more other regions. Specifically, both right and left putamen were correlated with each other, which is expected during monetary reward processing [14]. Further, both clusters of the right caudate (clusters 7 and 8) were highly correlated with each other and with the right putamen (cluster 1), which is supported by the previous findings of bidirectional anatomical connectivity between the putamen and caudate nuclei [14,63], suggesting dynamic interplay between the caudate nucleus and putamen in reward-related, instrumental behaviors [64]. Further, the right superior parietal lobule (cluster 3) activation is correlated with that of left putamen (cluster 2) and right angular gyrus (cluster 4), possibly suggesting functional connectivity across these regions while processing potential monetary rewards [65]. Similarly, the right angular gyrus (cluster 4; BA 39) was correlated with the right inferior parietal lobule (cluster 10; BA 40), indicating a strong functional link across these areas of multimodal regions responsible for visuospatial attention and other higher cognitive functions during visual tasks involving stimulus evaluation [58]. Lastly, the correlation between the right rolandic operculum (cluster 6) and right supramarginal gyrus (cluster 9), which may represent an evaluation of the subjective emotional state associated with gambling outcomes, as these adjacent cortical regions are often related to subjective evaluation of emotions [61] and the affective states [66], respectively.

### 4.2. Associations Between the Reward Regions and Behavioral Features

Exploratory bivariate correlations identified a few important and meaningful associations across the individual variables with the r-values ranging from 0.3617 to 0.5603, suggesting moderate effect sizes (**Table 4**). Further, these associations are relevant and meaningful and may guide future studies to examine them systematically. Therefore, it is worth discussing these associations in a broader context.

#### 4.2.1. Associations Between the Reward Regions and Impulsivity

Exploratory bivariate correlations identified a few negative correlations between impulsivity and fMRI activation clusters (**Table 4**), which include (i) non-planning impulsivity with all four striatal structures such as the bilateral putamen and right caudate (clusters 1, 2, 7, and 8), (ii) motor impulsivity with right superior parietal lobule (cluster 3), and (iii) total impulsivity with right superior parietal lobule (cluster 3) and with right caudate (cluster 8). These findings indicate that higher impulsivity was associated with lower activation in those regions for the contrast of win-loss. In other words, those with heightened impulsivity showed either lower activation during the win processing or higher activation during the loss condition, and vice-versa may be true for those with lower impulsivity. While the theories of choice and decision-making posit that loss looms larger than gain in most individuals [67], our findings indicate that sensitivity to loss is reflected more in those with higher impulsivity. Specifically, all four clusters representing core reward structures of the striatum were correlated with this non-planning impulsivity, a predisposition toward rapid, unplanned reactions to internal or external stimuli without regard to the negative consequences [25], suggesting that impulsive people manifest an urge for immediate gratification, regardless of whether the immediate reward is certain or uncertain [21]. On the other hand, motor impulsivity and total impulsivity showed a negative association with the right superior parietal lobule, a region that is modulated by the reward amount and its probability while performing decision-making tasks [68]. Although the total impulsivity score was also correlated with the right caudate (cluster 8), this was mostly contributed by the non-planning score. Interestingly, there were no significant correlations of age and education with any of the fMRI activation clusters.

#### 4.2.2. Associations Between the Reward Regions and Gambling Performance

With regard to associations between performance variables of the gambling task and the fMRI activation clusters, the exploratory analysis identified a few meaningful correlations (**Table 4**). Interestingly, the right putamen (cluster 1) showed (i) positive correlations with risky choice such as betting with 50 (bigger amount) following a loss during previous trial and previous two trials of the gambling task and (ii) negative correlations with safer choice such as betting with 10 (smaller amount) following a loss during previous two and three trials. This finding of higher activation of right putamen for risky bets and lower activation for safer or less risky bets suggests that right putamen is modulated by how much is at stake and how much risk or loss is anticipated (i.e., loss sensitivity) during each trial. This finding is consistent with that of other studies that putamen was associated with the stimulus-action-reward association [28] and low putamen activity was associated with poor reward sensitivity [29]. Further, the risky choices (i.e., higher bets in the face of previous loss) were also positively correlated with the right rolandic operculum (cluster 6), a cortical region associated with negative affective states such as apathy, depression, anxiety, and perceived stress [61], possibly representing a stressful state of anticipating a potential negative outcome during the gambling task. While our findings are very meaningful in the context of neural correlates underlying reward processing during a monetary gambling task, more studies with larger sample sizes are needed to further confirm and explain these preliminary findings.

#### 4.2.3. Associations Between the Reward Regions and Neuropsychological Scores

The results from the current study also pointed at a couple of associations between neuropsychological variables and the fMRI activation clusters (**Table 4**). First, a negative correlation was observed between right putamen activation and the total correct score of the total items correctly remembered during the visual span test representing short-term memory capacity. This may indicate that individuals who had poor short-term memory capacity showed relatively higher activation during the win condition (compared to the loss condition), or conversely, those with higher short-term memory capacity showed lower activation for the win condition (relative to the loss condition). In other words, putamen activation during reward processing varied based on the short-term memory capacity of the individuals. A previous study reported putamen may modulate working memory [31] during a delayed-response task that required memory updating. Putamen was also found to modulate cognitive functions in general [32] and learning and memory in particular [33,34]. It is possible that basal ganglia and medial temporal lobe memory systems work together in a complementary manner based on the task at hand [69]. However, more studies are needed to confirm the exact role of putamen under specific task conditions.

Further, one of the fMRI activation clusters, the left inferior occipital cortex, was negatively correlated with excess moves made (i.e., poor planning) during the Tower of London Test suggesting that higher activation of this brain region was associated with better cognitive planning. Although the inferior occipital cortex (Brodmann area 18), which is the secondary visual association cortex, does not have a specific role in reward processing per se, it is shown to be activated during visual-spatial tasks requiring visual attention to process and integrate various features of visual stimuli including color, shape, texture, and brightness [70] and to identify objects representing specific contexts [71]. While it is understandable that the Tower of London test, a visual task presented with initial and target positions of beads in different colors, would require a strong involvement of the inferior occipital cortex during task performance, the association of its activation level with cognitive planning ability is a novel finding, which may require further empirical support and interpretation.

#### 4.2.4. Clinical Implications

Empirical evidence supports reward-network dysfunction in substance-use disorders [72] and other psychiatric disorders [73]. Problems with both impulsivity and reward processing underlie several psychiatric disorders [74], including substance use disorders [75–80], attention-deficit hyperactivity disorder [81,82], antisocial personality [83,84], conduct disorder [85,86], borderline personality disorder [87,88], eating disorder [89,90], and gambling addiction [91,92]. Therefore, elucidating specific brain regions activated during various aspects of reward processing is essential to understanding, diagnosing, and treating these disorders [4,7,93], as specific abnormalities in reward processing can be observed in different forms of psychopathology [4]. Further, elucidation of reward dysfunction across a range of diagnostic categories may help to refine the phenotypes of brain structure and function and thus improve the prediction of onset and recovery of these disorders [94]. Furthermore, a better understanding of disorder-specific and/or symptoms-specific neural correlates of reward processing will help refine brain-based treatment techniques, such as brain stimulation and neurofeedback, in the management of substance use disorders and other related psychiatric disorders [95–97]. Lastly, elucidation of individualized brain connectome based individual’s symptom profile will help optimize personalized medicine approaches to treating a range of reward-related disorders [98].

#### 4.2.5. Limitations and Suggestions

While our study has produced some interesting findings, it has a few limitations. First of all, the sample size is relatively small, limiting the power needed to elicit associations between the clusters and the behavioral features. Second, the sample consisted of only males, and therefore the results are not generalizable to both genders. Third, the age range (19-38) is wide, and brain development and behavioral characteristics may vary between those in their 20s and 30s. Finally, the current study has only analyzed the win-loss contrast; other relevant contrasts (e.g., larger vs. smaller rewards, risky vs. safe bets, etc.) may also be important aspects of reward processing. We suggest conducting future studies with larger sample sizes consisting of both males and females to enhance the statistical power and generalizability of the findings. Future studies of a similar gambling task should analyze different age cohorts to examine reward processing across different developmental stages. Future studies may also consider examining other contrasts and paradigms of reward processing, while also examining different measures (e.g., functional connectivity).

## 5. Conclusions

The current study was designed to elicit neural substrates underlying reward evaluations of win versus loss outcomes in a monetary gambling paradigm as well as to understand possible associations of these brain regions with behavioral characteristics such as impulsivity, task performance, and neuropsychological measures. Findings revealed that a set of key brain structures, such as putamen, caudate nucleus, superior and inferior parietal lobule, angular gyrus, and Rolandic operculum, showed greater activation during win relative to loss condition and most of them were highly correlated with each other. Although the multivariate canonical correlation analyses failed to elicit associations between the anatomical regions and behavioral characteristics, exploratory bivariate correlations were significant with moderate effect sizes. Additionally, some of these reward-related regions showed meaningful associations with specific features of impulsivity, task performance, and neuropsychological measures. Further studies with larger samples are needed to confirm these preliminary findings.

